# Pre-saccadic remapping relies on dynamics of spatial attention

**DOI:** 10.1101/293886

**Authors:** Martin Szinte, Donatas Jonikaitis, Dragan Rangelov, Heiner Deubel

## Abstract

Each eye movement shifts the projections of the visual scene on the retina. It has been proposed that the receptive fields of neurons in oculomotor areas are remapped pre-saccadically to account for these shifts. While remapping of the whole visual scene seems prohibitively complex, selection by visual attention may limit these processes to a subset of attended locations. Because attentional selection consumes time, remapping of attended locations should evolve in time, too. In our study, we cued a spatial location by presenting an attention capturing cue at different times before a saccade and constructed detailed maps of attentional allocation across the visual field. We observed no remapping when the cue appeared shortly before saccade. In contrast, when the cue appeared sufficiently early before saccade, attentional resources were reallocated to the remapped location. Our results suggest that pre-saccadic remapping is an attentional process relying on the spatial and temporal dynamics of visual attention.

## Introduction

Our eye movements shift the visual scene on our retinas. Nevertheless, these shifts go largely unnoticed and don’t prevent us to efficiently interact with objects surrounding us. It has been proposed that the visual system compensates for these shifts using a copy of the motor command^1^ to anticipate changes in the visual scene due to the planned eye movement. Such an active mechanism could maintain an impression of space constancy and allow us to effectively interact with visual objects. However, we typically do not keep track of the whole visual scene^2,3^. Later studies proposed that such visual compensation could be restricted to salient or task relevant objects, selected by spatial attention^4,5^. At the behavioral level, this compensation could result in anticipatory deployment of spatial attention to the retinal location a visual stimulus will occupy after the saccade^6-9^. Such anticipatory deployment could explain observations that attention is allocated at a spatial target location almost immediately after a saccade^7,10^.

At the neuronal level, these visual compensations have been described as a remapping of visual neuron receptive fields. Remapping triggers an anticipatory and, sometimes, pre-saccadic response of neurons in Frontal Eye Fields (FEF), Lateral Intra-Parietal area (LIP) and Superior Colliculus (SC) with receptive fields centered on the post-saccadic retinal location of the attended object^11-13^. Remapping could facilitate tracking of task relevant objects across saccades and allow a rapid comparison between pre- and post-saccadic visual inputs^14^. However, this remapping hypothesis has been challenged with new data collected within the FEF^15^. In this study authors showed that, before a saccade, neurons respond to stimuli presented near the saccade target rather than to stimuli presented at remapped locations of the recorded RF. These results were later termed “convergent remapping” towards the saccade target in dissociation of the “forward remapping”, which would be parallel to the saccade vector^16^. They led to the proposal that convergent remapping could manifest behaviorally as a spatially unspecific spread of attention around the saccade target^17^. Remapping of spatial attention before saccades, as reported in behavioral studies, therefore could be reinterpreted as attentional spread between saccade target and remapped location^6-9^. Such interpretation of the convergent remapping effects predicts that locations surrounding the saccade target by up to 10 degrees of visual angle would receive all attentional benefits before the eyes start to move. In contrast, the forward remapping hypothesis predicts distinct pre-saccadic focuses of attention, separately for the saccade location and other attended locations. Critically, all attended locations would be remapped in the direction parallel to the saccade vector. Up to date, there is no behavioral study that mapped the pre-saccadic attention in sufficient detail to disambiguate these hypotheses.

We developed a protocol that allowed us to measure detailed maps of pre-saccadic attention, by measuring the orientation sensitivity at multiple locations while participants prepared a saccade (Figure 1). We observed that attention was allocated to the saccade target location and did not spread around it. Next, we measured remapping of attention in the presence of a salient cue during a saccade task. Contrary to neurophysiology studies, that typically presented the visual distractor just before eye movement onset, we manipulated the timing of stimulus onset relative to the saccade time. Our reasoning was that if remapping is an attentional process^4,5^, it will take some time for the attentional shift to occur^18-21^. Therefore, stimuli presented just before a saccade would not leave enough time for remapping to develop. On the other hand, stimuli presented early enough should be remapped before the saccade. Indeed, we found that when the cue appeared shortly before saccade onset, spatial attention was allocated at the cued location but not at its remapped location. In contrast, when the cue appeared sufficiently early before saccade onset, attentional resources that were initially drawn to the cued location were now reallocated to its remapped location (i.e. the retinal location it will occupy after the saccade).

**Figure 1.**
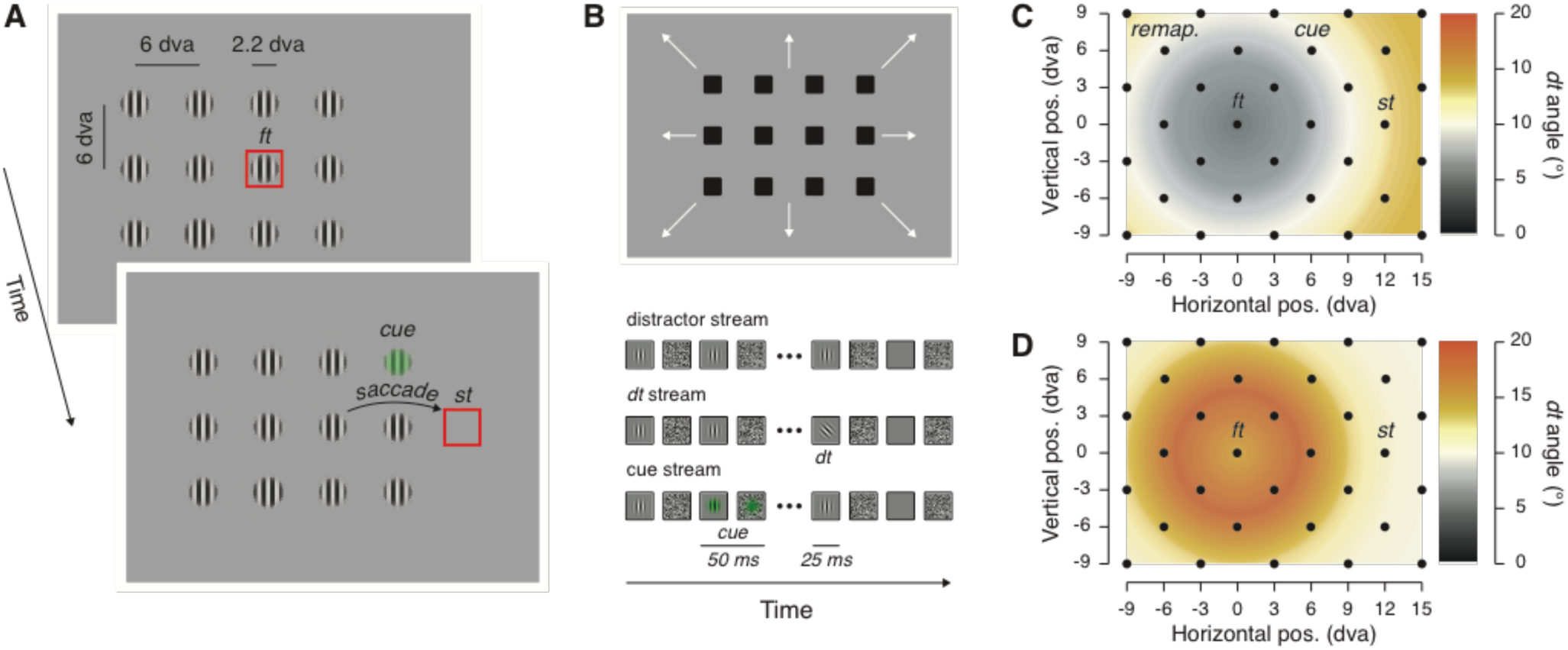
Stimuli displays and stimulus eccentricity effects. **A.** Participants fixated on the fixation target (*ft*) and prepared a saccade towards the saccade target (*st*). On each trial 12 visual streams (40 Hz flickering vertical Gabors and masks) were shown and in 2/3 of the trials a cue was flashed (50 ms) either above or below a virtual line between the fixation and the saccade targets (note that the stimuli are here sketched to increase their visibility, actual stimuli match those shown in the visual stream depiction). **B.** The arrangement of visual streams could take several positions (see Methods), such as to cover the whole display across trials. Participants reported the orientation of a discrimination target (*dt*), a tilted Gabor. **C-D.** The discrimination target was shown across trials at 32 different positions (see black dots) covering 24 degrees of visual angle (dva) horizontally and 18 dva vertically (see black dots) and including 4 main position of interest (the fixation target: *ft*; the saccade target: *st*, the cue: *cue*; and the remapped location of the cue: *remap.*). The tilt angle of the discrimination target was titrated to yield comparable performance at differently cued eccentricities from the fixation target. We adjusted these tilts in a preliminary task made either while participants keep their eyes steady at the fixation target (**C**: peripheral remapping threshold task in *Methods*) or prepared a saccade (**D**: foveal remapping threshold task in *Methods*). The maps show *dt* tilt angles averaged across participants in these two threshold tasks.

## Results

We measured detailed maps of spatial attention before a saccade in two different conditions: first, when participants made a visually guided saccade, and second, when a transient peripheral stimulus, a cue, was additionally presented during the preparation of a visually guided saccade. We assessed spatial attention by asking participants to report the orientation of a briefly presented tilted discrimination target (clockwise or counterclockwise tilted Gabor), embedded in a display of vertical distractor streams (vertical Gabors, Figure 1A-B). To ensure that the discrimination task could be solved correctly only if participants attended at a particular location, we first completed a threshold task in which participants fixated at the center of the screen. This threshold task was used to estimate the tilt angle of a cued discrimination target presented at different eccentricities from the fixation. We observed that in order to achieve a comparable discrimination at different eccentricities, the discrimination target had to be tilted by 4.42 ± 0.86° (mean ± SEM), if presented at the fixation target. This tilt gradually increased with eccentricity, finally reaching 14.1 ± 1.4° at an eccentricity between ~15.3 and ~16.2 dva (see Figure 1C). We used these threshold tilt values at their respective eccentricities in the main saccade task. This ensured that discrimination performance reflected the modulation of spatial attention over space rather than visual acuity.

We first verified that the presentation of the discrimination target during saccade preparation did not disrupt eye movements. Such a disruption, as measured by saccade latency or accuracy, would suggest that the stimuli used to measure attention instead captured attention. For this we first determined whether the eccentricity of the discrimination target affected saccade latency. Saccade latencies were longer when the visual streams overlapped with the fixation and saccade targets (217.56 ± 3.77 ms) as compared to when they didn’t overlap (186.00 ± 2.61 ms, *p* < 0.0001). This indicated that such difference was due to the saccade target and fixation being less visible if they overlapped with the visual streams. Therefore, we separated the trials based on whether the fixation and saccade targets overlapped with the visual streams or not. Discrimination target eccentricity did not affect saccade latency on trials in which the fixation and saccade targets overlapped with visual streams (repeated measures ANOVA with discrimination target eccentricity as the main factor, F_3,39_ = 0.08, *p* > 0.05) or on trials when they didn’t (F_3,39_ = 1.49, *p* > 0.05). Note that from a pilot study, we expected to find such saccade latency cost during the above-mentioned trials, and correspondingly presented the discrimination target 25 ms (1 visual stream) later on those trials. This procedure ensured a homogenous timing of the discrimination target relative to the saccade onset irrespectively of the tested position. Next, we evaluated whether discrimination target eccentricity affected saccade accuracy (i.e. the absolute distance between the saccade target and the saccade landing point), as would be evident if the target captured attention. Again, we didn’t find such an effect (main effect of discrimination target eccentricity, F_4,52_ = 2.11, *p* > 0.05). Altogether, these results show that the onset of the discrimination target did not disrupt the saccade preparation, demonstrating that the stimuli used to measure the allocation of attention did not directly interfere with its deployment.

We next measured the pre-saccadic allocation of attention. Trials with and without a cue were analyzed separately. On trials without a cue (Figure 2A-C), we found increased visual sensitivity (0.88 ± 0.05, normalized d’ and SEM, respectively) at the saccade target location relative to the average of all other tested positions (0.42 ± 0.04, *p* < 0.0001, Figure 2B) suggesting that attention shifts towards the saccade target during the saccade preparation. Further, we tested the spatial specificity of this effect by comparing visual sensitivity at the saccade target location with the averaged visual sensitivity at the four positions surrounding it. Attention at the saccade target clearly did not spread to the surrounding positions (0.40 ± 0.04, *p* < 0.0001, Figure 2C), with sensitivity benefits being constrained within ~4.2 dva around the saccade target. Also, we did not observe a deployment of spatial attention to the fixation target (0.41 ± 0.05) as compared to the average across all other positions (*p* > 0.05).

**Figure 2.**
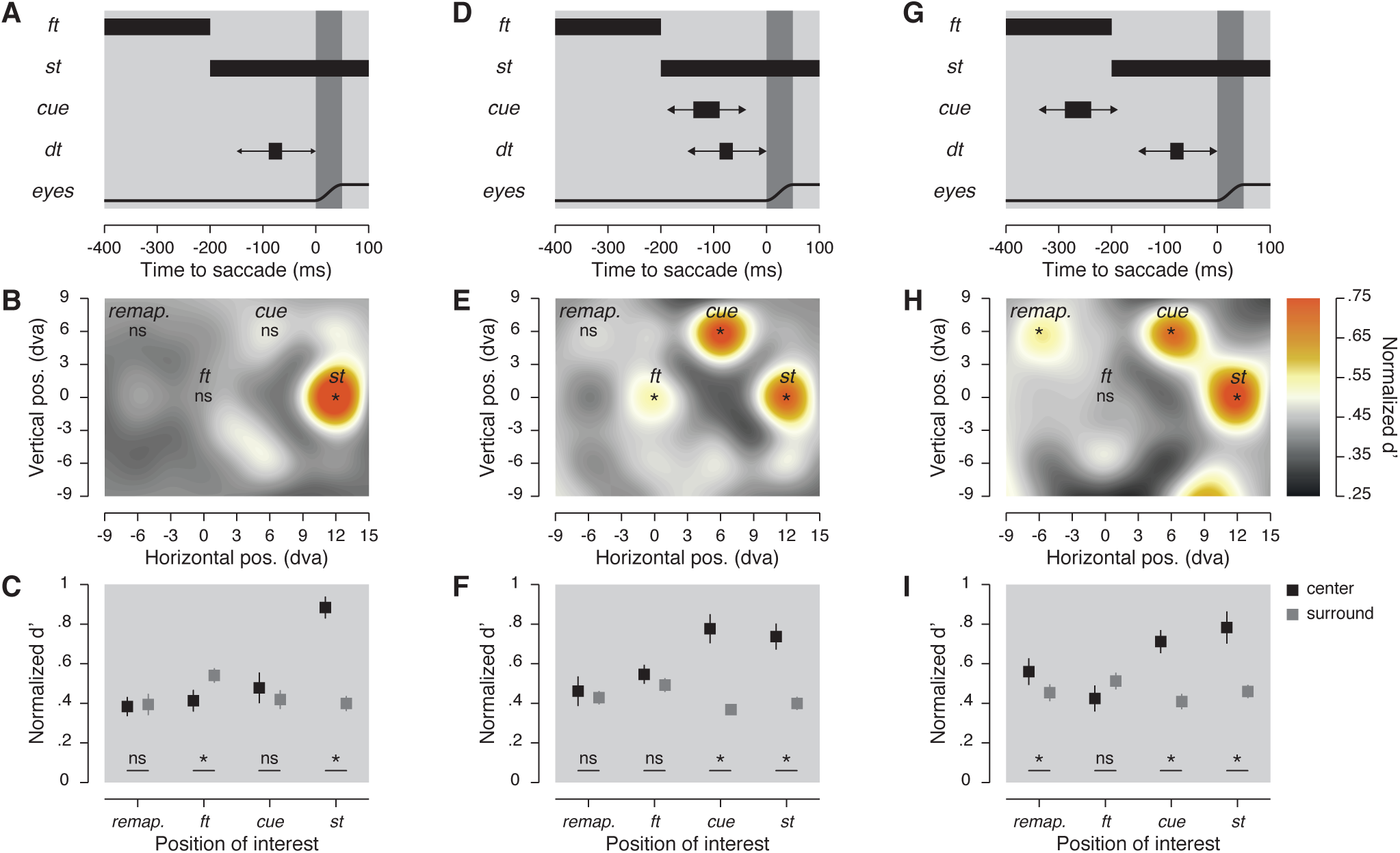
Stimulus timing and sensitivity maps. **A-D-G**. Stimulus timing. Participants prepared a saccade at the offset of the fixation target (*ft*) which corresponded to the onset of the saccade target (*st*). In the last 150 ms before the saccade a discrimination target (*dt*) was briefly shown at one of the 32 possible positions. A cue was then either never shown (**A**), shown 50 ms before the *dt* and about 100 ms before the saccade (**D**), or shown 200 ms before the *dt* and about 250 ms before the saccade (G). **B-E-H**. Normalized sensitivity maps. Averaged normalized sensitivity (d’) observed across participants and displayed using a color-coded linear scale going between 0.25 and 0.75 (see Methods). Asterisks indicate significant differences (*p* < 0.05) in sensitivity found between a particular position of the *dt* and the average of all the others tested positions. **C-F-I.** Averaged normalized d’ obtained at four positions of interest (black squares) and at their corresponding surrounding positions (dark gray squares). Error bars show SEM and asterisks indicate significant comparisons (*p* < 0.05).

We next analyzed the trials during which we presented an additional cue during saccade preparation, with the cue and discrimination target shown shortly after each other (50 ms) and on average 96.88 ± 0.96 ms (cue offset relative to saccade onset) before the saccade (Figure 2D-F). Here, in addition to the fixation and saccade targets, we were also interested in two further locations: the cue and the remapped location of the cue (the retinal location the cue will occupy after the saccade). Visual sensitivity was higher at the cue (0.78 ± 0.07, *p* = 0.0004), at the saccade target (0.74 ± 0.07, *p* < 0.0001) and at the fixation target (0.55 ± 0.05, *p* = 0.0038), when compared to the average of all the tested locations (0.44 ± 0.03). When compared to their closest surrounding positions (Figure 2F), we found spatially specific effects only at the cue (surround: 0.37 ± 0.03, *p* < 0.0001) and at the saccade target (surround: 0.40 ± 0.03, *p* < 0.0001), but not at the fixation target (surround: 0.49 ± 0.03, *p* > 0.05). Importantly, when the cue was shown shortly before the saccade, the visual sensitivity at its remapped location (0.46 ± 0.07) was not significantly higher relative to other tested positions (0.44 ± 0.03, *p* > 0.05). Thus, we observed no evidence for forward remapping of the cued location when the cue appeared shortly before saccade onset.

These results contrasted with those found on trials when the same cue was shown substantially before the discrimination target (200 ms) and on average 240.82 ± 1.42 ms before the saccade onset (Figure 2G-I). Under such conditions, when compared with the average across all positions (0.45 ± 0.08), we found higher visual sensitivity at the saccade target (0.78 ± 0.08, *p* < 0.0001), at the cue (0.71 ± 0.06 *p* < 0.0001) and, critically, at the remapped location of the cue (0.56 ± 0.07, *p* = 0.0068). Similar to other experimental conditions, the benefits observed at the saccade target (surround: 0.46 ± 0.03, *p* < 0.0010), at the cue (surround: 0.41 ± 0.04, *p* < 0.0001) and at its remapped location (surround: 0.45 ± 0.04, *p* = 0.0352) did not spread towards their respective adjacent positions. Combined, our results so far show that when the visual system is given enough time to process and attend a visual stimulus, such as the salient cue used in our task, spatial attention is remapped to a retinal location the stimulus will occupy after a saccade.

If attention is remapped pre-saccadically, one should also expect to find spatially specific attentional effects at the fixation target, as the fixation target is the remapped location of the saccade target. Indeed, one study reports such foveal remapping of the saccade target^6^. In the experiment above, we observed inconsistent evidence for spatial attention at the fixation target (see Figure 2C, 2F, 2I). This is likely due to the threshold procedure differences between this study and the one that reported spatial remapping of attention to fixation^6^. Indeed, as we were principally interested in remapping of the salient peripheral cue in above study, our threshold procedure measured spatial attention during fixation (giving, importantly, no threshold difference between the cue and the remapped location of the cue). However, previous research has shown that preparation of saccades draws spatial attention away from other, non-saccade or non-salient locations^22^. Since in the experiment described above, we measured perceptual thresholds in a fixation task, rather than in a saccade task, it is possible that the threshold for the fixation location underestimated the one during saccadic preparation. If it was the case, potential remapping of the saccade location to the fixation location could have been masked by drawing attention away from the fixation location in the threshold task. Therefore, in the threshold task of a second experiment (foveal remapping task) we adjusted the tilt of the discrimination target using a saccade rather than a fixation task (see foveal remapping main task in *Methods*). We observed that the discrimination target had to be tilted by 15.03 ± 2.34° if presented at the fixation target before a saccade (Figure 1D), a tilt up to four times bigger than that recorded during the fixation threshold procedure (Figure 1C). We used these values in a simplified version of the above experiment, without the presentation of the cue, and as before, with a discrimination target randomly shown across trials at 32 possible positions while participants prepared a visually guided saccade. If anything, this procedure should be even more sensitive at detecting attention spread between saccade and fixation targets^17^.

Then in this second task, in which only such thresholding procedure changed, we first analyzed whether the discrimination target affects saccade latency and accuracy. We found a main effect of the eccentricity of the discrimination target on saccade latency within trials in which the visual streams overlapped with the fixation and the saccade targets (F_3,21_ = 8.89, *p* = 0.0005), but not on trials when the visual streams did not overlap with the targets (F_3,21_ = 3.01, *p* > 0.05). Further, discrimination target eccentricity did not affect saccade accuracy (F_4,28_ = 1.60, *p* > 0.05). These results suggest that, similar to the first experiment, the presentation of the discrimination target had limited influence on the preparation of the saccade. Next, and in agreement with the results of the first experiment, we found a systematic pre-saccadic deployment of attention towards the saccade target (0.68 ± 0.11, *p* = 0.0004) when compared to the average over all the tested positions (0.36 ± 0.03). Critically, this benefit was accompanied by a systematic deployment of attention at the fixation target (0.96 ± 0.02, *p* < 0.0001). Finally, these effects were spatially specific (Figure 3B), as shown by the significant differences observed when comparing sensitivity at the fixation target (surround: 0.47 ± 0.06, *p* < 0.0001) and the saccade target (surround: 0.33 ± 0.04, *p* < 0.0001) to their relative surrounds.

**Figure 3.**
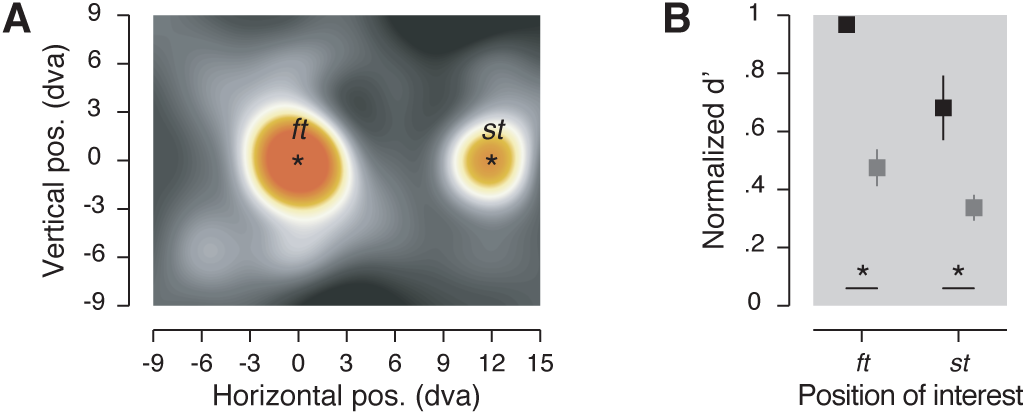
Foveal remapping task results. **A.** Normalized sensitivity maps. Averaged normalized sensitivity (d’). **B.** Averaged normalized sensitivity (d’) obtained at two positions of interest (see center in black) and at their corresponding surround positions (see surround in dark gray). Conventions and color scale are as in Figure 2.

## Discussion

We constructed detailed spatial attention maps by measuring orientation sensitivity while participants prepared a visually guided saccade. Our paradigm allowed us to measure whether attention broadly spread around the saccade target or shifted towards spatially specific loci. We observed that attention consistently shifted to the saccade target location and, importantly, did not spread to other locations surrounding it. In our main manipulation, we presented a salient cue during saccade preparation. On these trials, we observed a second spatially specific locus of attention at the cued location. Importantly, on these cued trials, we observed a third, distinct locus of attention. Although this third location was neither salient, nor task relevant before the saccade, it corresponded to the retinotopic location the cue will occupy after the saccade. In other words, we observed remapping of the cue location before the saccade onset, an effect consistent with the forward remapping hypothesis. Critically, we observed these effects exclusively when the cue appeared long enough before the saccade onset. This indicates that forward remapping, like any other attentional process, requires some time. As we observed three separate focuses of attention (at the saccade target, cue and remapped location), we definitely ruled out the hypothesis that during saccade preparation attention spreads around the saccade target, as expected by the convergent remapping hypothesis^15,23,24^.

Our findings speak to the current debate on whether forward remapping exists and what role it plays in maintaining visual stability. Behavioral studies on visual remapping of attention^6-10^ have been inspired by earlier neurophysiology work suggesting that receptive fields in FEF, LIP and SC shift (or are remapped) in anticipation of post-saccadic stimulus location^11-13^. Such predictive receptive field shift occurs either shortly after a saccade (at a neural latency too short for visual responses) or even before a saccade onset^25,26^. Forward remapping offered an excellent candidate mechanism of space constancy which was incorporated into such phenomenon models. In these models, connections between visual neurons with receptive fields spatially separated by the saccade vector can be used to predict post-saccadic visual stimulus location and compensate for retinal image shifts during the saccade^27-29^. However, forward remapping models have been challenged by Zirnsak and colleagues^15^, whose results suggested that the early findings have been affected by the low spatial resolution of receptive field mapping technique^11-13^. A more detailed FEF receptive field mapping suggested that, before a saccade, cells preferentially respond to stimuli presented nearby the saccade target rather than at the remapped target location. In other words, cell receptive fields did not shift in parallel to the saccade vector, but instead converged towards the saccade target. Zirnsak and colleagues thus argued that forward remapping models can’t explain space constancy and instead one should focus on saccade target selection as a mechanism mediating this phenomenon^17^. A more recent work indicated that the visual system may, in fact, implement both forward and convergent remapping of receptive fields in area V4^16,30^. As this combined approach has been criticized on technical grounds^23^, the neurophysiological results regarding the existence of forward remapping remain inconclusive.

On the other hand, a number of behavioral studies have repeatedly and reliably demonstrated forward remapping of spatial attention before saccades^6-10^. The behavioral studies, however, suffered from the same drawbacks as early neurophysiological studies, a low spatial resolution. Our study has eliminated this potential criticism, and our results provide an unequivocal support for the forward remapping effects. Consistent with previous behavioral studies^31-33^ and contrary to the convergent remapping effects, our results show that spatial attention is allocated to the saccade target and does not spread around it. Additionally, convergent remapping cannot account for a number of earlier behavioral findings^6-9^, as such spread of attention would have to be asymmetric and not spread towards the several control positions tested in these earlier studies.

Of note, the behavioral consequences of pre-saccadic changes in the spatial tuning of visual cells RF are unclear. Perhaps counterintuitively, recent computational neuroimaging modeling^34^ has shown that increasing the neural spatial sampling at a particular position, similar as the over-sampling of the saccade target observed within convergent remapping studies^15,23,24^, results in a reduction of spatial uncertainty. Convergent remapping then does not necessarily yield a large spread of attention around the saccade goal^17^. Rather, it may increase visual sensitivity to stimuli only in the immediate vicinity of the saccade target. Convergent remapping could, therefore, reflect the spatially specific attentional selection of the saccade target observed in the present study. Our results obtained with a peripheral cue are, however, not consistent with convergent remapping effects. To account for our attentional remapping effects, one must assume a mechanism similar to forward remapping in which attended locations are remapped in the direction parallel to the saccade vector. Our results indicate that spatial visual attention mechanisms must be accounted for in future work of remapping in order to advance our understanding of space constancy. Here, we hypothesize that previous reports of convergent remapping likely reflect increased visual sensitivity at the saccade target explained by both spatial and temporal properties of visual attention.

First, to determine a visual neuron RF spatial profile, neurophysiologists used localized visual probes, presented most of the time in a sparse display with high probe-background contrast. It is, therefore, likely that such probes capture spatial attention^35^. The same holds for visual stimuli used to trigger the saccade, which were both task-relevant and, in most experiments, high-contrast stimuli. As visual RFs can shift towards attended locations even without any saccade involved^36,37^, one must account for the effect of attention before interpreting any RF change in spatial tuning. In our study, the attention-capturing cue and the saccade target were dissociated from the measure of spatial attention. We measured attention by a discrimination target which did not significantly capture attention on its own, and, therefore, did not interfere with saccade preparation or pre-saccadic distribution of spatial attention^33^. We also use different conditions to account separately for the effect of the saccade target and of the cue. In order to understand pre-saccadic RF changes in spatial tuning, one should first consider the spatial deployment of spatial attention.

Second, we argue that the temporal dynamics of attention should also be accounted for. We observed that the time at which we presented our cue was critical for forward remapping to occur. In particular, benefits at the remapped location of the cue were observed only when the cue was shown more than 175 ms before the saccade onset. As even the fastest deployment of attention would take a minimum of 100 ms to occur^18-21^, our remapping effects are well compatible with the time course of attentional selection. It is important to note that convergent remapping, in contrast, was typically observed when probes were shown within the last 50 ms preceding the saccade^15,16,23^. Our findings therefore suggest that the short time window between probe presentation and neural recording, which these studies have used, was insufficient for attention to be remapped at a location parallel to the saccade. Further, if remapping is closely related to the time course of attention, it is possible that for attended stimuli shown just prior to the saccade onset, remapping may occur during or after the saccade. Indeed, Neupane and colleagues^16,30^ observed forward remapping when measuring post-saccadic memory responses to probes shown just before the saccade. Also, Yao and colleagues^38^ have shown that a post-saccadic memory trace of remapping was enhanced by attentional modulations established before the saccade, corroborating the notion that forward remapping can occur after the saccade. Although we did not measure whether a cue shown shortly before saccade onset was remapped after the saccade in the present study, two earlier studies did^7,10^. Both studies found that spatial attention was allocated to the location of a salient stimulus immediately after the saccade, even when the stimulus was no longer present^7^. This indicates that the visual system anticipated the attended stimulus location after a saccade and recomputed its retinotopic location before the saccade is done.

Finally, we also observed high perceptual performance at fixation, a result in line with two earlier studies that investigated foveal remapping effects^6,39^. Such an attentional effect is surprising, as one would expect that visual selection should prioritize the saccade target, whereas the current fixation should be the least informative and least attended part of the display (especially given that participants already fixated for ~1 second before starting saccade preparation). However, if one considers that fixation-centered receptive fields will process the saccade target after the saccade, forward remapping effects would suggest significant attentional benefits at that location, as we observed here and in a previous study^6^.

In summary, we used an eccentricity-adjusted discrimination task to measure, for the first time, detailed spatial attention maps before saccades. Using this method, we observed a spatially specific increase in visual sensitivity at the fixation target, the saccade target, the cue and the remapped location of the cue. These results indisputably reject the hypothesis that spatial attention spreads around the saccade target in a spatially unspecific way, as predicted by convergent remapping hypothesis. We found that, before a saccade, attention is deployed towards the saccade target as well as towards a cued location. Further, given the cue was presented sufficiently early before the saccade, we observed a deployment of attention to its remapped location, that is parallel and opposite to the saccade vector. While the benefit at that location is smaller as compared to the cue location, it reflects an ongoing process that facilitates spatial attention allocation after the saccade despite the retinotopic shifts induced by the eye movement.

## Methods

### Participants

Eighteen students (14 participants in the peripheral remapping task, 8 participants in the foveal remapping task, 4 participants did both tasks) of the LMU München participated in the experiment (ages 22-30, 10 females, 1 author), for a compensation of 10 Euros per hour of testing. All participants except the author were naive as to the purpose of the study and all had normal or corrected-to-normal vision. The experiments were undertaken with the understanding and written informed consent of all participants and were carried out in accordance with the Declaration of Helsinki. Experiments were designed according to the ethical requirements specified by the LMU München and an institutional review board ethics approval for experiments involving eye tracking.

### Setup

Participants sat in a quiet and dimly illuminated room, with their head positioned on a chin and forehead rest. The experiment was controlled by an Apple iMac Intel Core i5 computer (Cupertino, CA, USA). Manual responses were recorded via a standard keyboard. The dominant eye’s gaze position was recorded and available online using an EyeLink 1000 Desktop Mounted (SR Research, Osgoode, Ontario, Canada) at a sampling rate of 1 kHz. The experimental software controlling the display, the response collection as well as the eye tracking was implemented in Matlab (MathWorks, Natick, MA, USA), using the Psychophysics^40,41^ and EyeLink toolboxes^42^. Stimuli were presented at a viewing distance of 60 cm, on a 21-in gamma-linearized Sony GDM-F500R CRT screen (Tokyo, Japan) with a spatial resolution of 1,024 x 768 pixels and a vertical refresh rate of 120 Hz.

### Procedure

We completed two different tasks (peripheral remapping and foveal remapping) in a total of 4 experimental sessions (on different days) of about 100 minutes each (including breaks). Each task was always preceded by a discrimination threshold measurement, completed at the beginning of each session. Each session was composed of 2 blocks of the threshold task followed by 4 to 6 blocks of the main task. Participants ran a total of 11-12 blocks of the peripheral remapping task and 4 blocks of the foveal remapping task. Participants who completed the two tasks always started with the peripheral remapping task.

### Peripheral remapping task

Each trial began with participants fixating a fixation target, a red frame (2.2 dva/side, 10’ width, 30 cd/m^2^) presented on a gray background (60 cd/m^2^). When the participant’s gaze was detected within a 2 dva radius virtual circle centered on the fixation target for at least 200 ms, the trial began with a random fixation period of 500-900 ms (uniform distribution, in steps of 50 ms). After this period, the fixation target was replaced by the saccade target (same red frame) presented 12 dva to the right or to the left of the fixation target (Figure 1A). Participants were instructed to move their eyes as quickly and as accurately as possible towards the center of the saccade target. From the beginning of the trial, we presented 12 flickering visual streams (40 Hz), composed of 25 ms vertical Gabor patches (frequency: 2.5 cycles per degree; 100% contrast; same random phase on each screen refresh; standard deviation of the Gaussian window: 0.9 dva; mean luminance: 60 cd/m^2^) alternating with 25 ms pixel noise square masks (2.2 dva side, made of ~0.04 dva-width pixels). The visual streams were arranged in 3 by 4 matrix, with a distance of 6 dva between each element (Figure 1A). On each trial the matrix of twelve visual streams was presented at 1 out of 15 different positions relative to the display center (shifted by -6 dva, -3 dva, 0, +3 dva or +6 dva vertically and -3 dva, 0 or +3 dva horizontally). Between 50 and 175 ms after the saccade target onset one of the 12 vertical Gabor patches was replaced by a discrimination target, a tilted Gabor (clockwise or counterclockwise from the vertical). The time interval of 50-175 ms was determined in a pilot study with two criteria: i) that discrimination target offset occurred in the last 150 ms before the saccade, and ii) taking into account shorter saccade latency on trials when the fixation and saccade targets were not covered by visual streams. Once the discrimination target had appeared, no more vertical Gabor patches were presented and only noise masks altered with blank frames (Figure 1B). Across trials the discrimination target was shown at 32 positions covering 24 dva horizontally and 18 dva vertically (position located at every second intersection of a 9 column by 7 rows grid, see Figure 1C-D). At the end of each trial, participants reported the orientation of the discrimination target using the keyboard (right or left arrow keys), followed by a negative-feedback sound on error trials.

In two thirds of the trials we captured attention by presenting a task-irrelevant cue, a 50 ms abrupt color onset stimulus presented in between the fixation and saccade targets, 6 dva above or below the screen center. This cue was a green Gaussian patch (mean luminance of 80 cd/m^2^), with the same Gaussian window of the Gabors and covering one of the visual streams. Across trials this cue was presented either 50 ms or 200 ms before the discrimination target onset. In one third of the trials the cue was not presented at all. To avoid inter-trial attention-lingering effect at the cue location, we separated cue and no-cue trials, with no-cue trials presented in the first four blocks of the task.

This method allowed us to map in detail the allocation of attention at 4 positions of interest: the fixation target, the saccade target, the cue, and the remapped position of the cue, as well as 28 control positions. To maximize the number of trials for statistical comparisons, we presented discrimination targets less often (30% less) in the two rows (9 positions) most peripheral from the cue (for example, if the cue was presented above the horizontal meridian, the discrimination target was presented less frequently at the two bottom rows below the horizontal meridian). In the trials without a cue, discrimination targets were presented less often (30% less) in the two most peripheral rows (18 top and bottom positions) maximizing the number trials around the fixation and the saccade target.

Participants completed between 2914 and 3323 trials of the peripheral remapping task. We checked correct fixation maintenance and correct saccade execution online and repeated incorrect trials at the end of each block. We also repeated trials during which a saccade started within the first 25 ms or ended after more than 350 ms following the saccade target onset (participants repeated between 159 and 443 trials).

### Peripheral remapping threshold task

On each session, before the peripheral remapping task, participants completed a threshold task. This allowed us to avoid possible effects of task learning across different sessions as well as to adjust the discrimination target tilt for different eccentricities from fixation. This latter point is particularly important as it reduced the impact of visual acuity^43^ on the measure of spatial attention. The threshold task was identical to the main task with the exception that participants kept fixation and the saccade target was not shown. After a random initial period of 500-900 ms (uniform distribution, in steps of 50 ms) a cue was briefly (50 ms) shown followed by a discrimination target 200 ms later at the cued location. Across trials the cue was shown at each of the 32 locations used in the main experiment. The positions of discrimination target and cue were subdivided into 5 equiprobable groups of eccentricity from the fixation target (eccentricity 1: the fixation target; eccentricity 2: from ~4.2 to 6 dva, eccentricity 3: from ~8.5 to ~9.5 dva; eccentricity 4: from 12 to ~13.4 dva and eccentricity 5: from ~15.3 to ~16.2 dva). We used a procedure of constant stimuli and a randomly selected orientation of the discrimination target from a linearly spaced interval for each eccentricity (each interval divided in 5 steps; between ±1 and ±9 dva for the first eccentricity, between ±1 and ±13 dva for eccentricity 2 and between ±1 and ±17 dva for eccentricities 3 to 5).

Participants were instructed that the cue indicated the position of the discrimination target and were told to report at the end of each trial its orientation (clockwise or counterclockwise). They completed 400 trials across 2 blocks of the threshold task. For each participant and experimental session individually, we determined for the five eccentricities from the fixation target, five threshold values corresponding to the discrimination target tilts leading to a correct discrimination in 85% of the trials. To do so, we fitted five cumulative Gaussian functions to performance gathered in the threshold blocks. These tilts were used in the main task at their respective eccentricity from the fixation target.

### Foveal remapping and threshold tasks

The foveal remapping task was identical to the peripheral remapping task with the exception that we did not present the cue. Moreover, the foveal remapping threshold task differed from the peripheral remapping threshold task, as participants made a saccade during the threshold task instead of keeping fixation^6^. In the foveal remapping threshold task the saccade target could be presented at any of the 32 locations tested. The discrimination target appeared 200 ms after the appearance of the saccade target and participants were instructed that the discrimination target could appear at either fixation or saccade target. We again used a procedure of constant stimuli, and chose discrimination target orientation randomly for intervals defined for the different eccentricities (intervals divided in 5 steps for each eccentricity; interval between ±1 and ±25 dva for the eccentricities 1 and 2 and between ±1 and ±21 dva for eccentricities 3 to 5).

Participants completed between 973 and 1137 trials of the foveal remapping main task. We checked correct fixation maintenance and correct saccade execution online and repeated incorrect trials at the end of each block. We also repeated trials during which a saccade started within the first 25 ms or ended after more than 350 ms following saccade target onset (participants repeated between 13 and 177 trials). Participants completed 500 trials across 2 blocks of the threshold task. For each participant and experimental session individually, we determined for the five eccentricities from the fixation target, five threshold values corresponding to the angels leading to correct orientation discrimination in 85% of the trials following the same procedure as in the peripheral threshold task.

### Data pre-processing

Recorded eye position data was processed offline (independent of online tracking during the experiment). Saccades were detected based on their velocity distribution^44^ using a moving average over twenty subsequent eye position samples. Saccade onset was detected when the velocity exceeded the median of the moving average by 3 SDs for at least 20 ms. We included trials if a correct fixation was maintained within a 2.0 dva radius centered on the fixation target, if a correct saccade started at the fixation target and landed within a 2.0 dva radius centered on the saccade target, and if no blink occurred during the trial. Finally, only trials in which the discrimination target disappeared in the last 150 ms preceding saccades were used in the analysis. In total, we included 36,236 trials (90.41% of the online accepted trials, 82.20% of all trials) in the peripheral remapping main task and 7306 trials (95.13% of the online accepted trials, 86.85% of all trials) of the foveal remapping main task.

### Behavioral data analysis

Data was analyzed separately for the 3 conditions of the peripheral remapping task and the only condition of the foveal remapping task. For the trials in which a cue was presented, the cue onset preceded the discrimination target onset by either 50 or 200 ms. Therefore, one condition included the trials with a SOA of 50 ms during which the cue disappeared in the last 175 ms before the saccade, and a second condition included the trials with a SOA of 200 ms during which the cue disappeared more than 175 ms before the saccade. A third condition of the peripheral remapping task and the all the trials of the foveal remapping task included trials in which no cue was shown.

To map the allocation of attention, we first mirrored discrimination target positions of leftward saccade trials to match those of the rightward saccade trials. Moreover, in trials with a cue, we mirrored positions of the bottom cue trials (trials in which the cue was shown 6 dva below the screen center) to match those of the top cue trials. Then, for each participant and each condition, we determined the sensitivity in discriminating the orientation of the discrimination target (d’): *d’ = z(hit rate) - z(false alarm rate)*. To do so, we defined a clockwise response to a clockwise discrimination target (arbitrarily) as a hit and a clockwise response to a counterclockwise discrimination target as a false alarm. Corrected performance of 99% and 1% were substituted if the observed proportion correct was equal to 100% or 0%, respectively. Performance values below the chance level (50% or d’ = 0) were transformed to negative d’ values. We next normalized for each participant individually, the sensitivity obtained at each position by the range obtained across all tested positions following this formula *d’_n_ = (d’_n_ - min) / (max - min)*, with *d’_n_* the sensitivity at a given *n* position, *min* and *max*, respectively the minimum and maximum sensitivity obtained across the 32 tested positions in the specific condition. These normalized values were then averaged across participants and used to plot sensitivity maps and to test statistical comparisons.

We obtained sensitivity maps (Figure 2B, 2E, 2H and 3A), by interpolating (triangulation-based natural neighbor interpolation) the missing values located at every second intersection of the 9 columns by 7 rows grid of discrimination targets using the 32 tested positions. We then rescaled the grid (Lanczos resampling method) to obtain a finer spatial grain. We drew position sensitivity maps across participants as colored maps coding the mean sensitivity across participants following a linear color scale going from 0.25 to 0.75 normalized sensitivity. To determine the threshold maps (Figure 1C-D), we first interpolated (linear interpolation) the mean threshold angle obtained for each participant individually over the 5 different distances between the fixation target and the discrimination target. Map of threshold *dt* angles across participants was then obtained by drawing disks centered on the fixation target with a radius corresponding to the eccentricity at which the discrimination target was played and coded the mean threshold angle obtained across participants following a linear color scale going from 0**°** to 20° of discrimination target tilt.

For statistical comparisons we drew (with replacement) 10,000 bootstrap samples from the original pair of compared values. We then calculated the difference of these bootstrapped samples and derived two-tailed *p* values from the distribution of these differences. Statistical comparisons of the eccentricity effect of the discrimination target on different saccade metrics (saccade latency and accuracy) was tested using repeated measures ANOVA. Discrimination target positions were grouped depending on their eccentricities from the fixation target as defined in the threshold tasks.

## Acknowledgments

This research was supported by a Deutsche Forschungsgemeinschaft temporary position for principal investigator grant to M.S. (SZ343/1) and D.R. (RA2191/1-1). We are grateful to the members of the Deubel laboratory in Munich for helpful comments and discussions and to Elodie Parison, Alice and Clémence Szinte for their invaluable support.

## Author contributions

M.S., D.J., D.R., and H.D. conception and design of research; M.S. performed experiments; M.S. analyzed data; M.S., D.J., D.R., and H.D. interpreted results of experiments; M.S. prepared figures; M.S. and D.J. drafted manuscript; M.S., D.J., D.R., and H.D. edited and revised manuscript; M.S., D.J., D.R., and H.D. approved final version of the manuscript.

